# Microbial community regulation of extracellular enzyme production can mediate patterns of particulate and mineral-associated organic matter accumulation in undersaturated soils

**DOI:** 10.1101/2025.07.25.666799

**Authors:** Paige M. Hansen, Yao Zhang, Ksenia Guseva, Christina Kaiser, M. Francesca Cotrufo

## Abstract

Dissolved low molecular weight (LMW) compounds in soil can either be assimilated by microbes or sorb onto mineral surfaces, forming mineral-associated organic matter (MAOM). This creates potential ‘competition’ between microbes and mineral surfaces for LMW compounds, potentially influencing whether particulate organic matter (POM) is retained or depolymerized by microbes to produce LMW substrates. Therefore, microscale interactions between unoccupied mineral surfaces and microbial enzymes may mediate patterns of POM and MAOM storage, particularly in soils varying in MAOM saturation. To explore this, we adapted an individual-based microscale model to simulate POM retention and new MAOM formation under different initial POM qualities (carbon:nitrogen ratio; C:N) and MAOM saturation levels, while also considering microbial social-like dynamics, which emerge from interactions between microbes with different capacities to produce and share public goods (in this case, extracellular enzymes). Consistent with prior findings, the presence of these dynamics slowed decomposition of initial POM pools. Additionally, MAOM saturation affected microbial community properties, new MAOM formation, and POM decomposition only when social dynamics were included. The patterns of POM decomposition and MAOM formation identified in our work align with observations of simultaneous POM and MAOM formation in undersaturated soils from prior field studies, suggesting that regulation of enzyme production via microbial interactions may be an additional driver of POM and MAOM storage in such soils. Overall, this highlights the importance of explicitly incorporating microbial ecology into our conceptual understanding of C and N cycling, particularly to improve the predictive capacity of ecosystem models and inform soil management strategies that enhance global change mitigation, especially in degraded soils likely to be undersaturated.

## 1. Introduction

Rising levels of atmospheric carbon dioxide (CO_2_) significantly impact the global climate, resulting in continuous warming and necessitating the development of innovative strategies to drawdown atmospheric CO_2_ while reducing CO_2_ emissions to the atmosphere. Soils, which represent the largest terrestrial carbon (C) store, may play a critical role in global change mitigation efforts through storage of atmospheric CO_2_ (Smith, 2016). In conjunction with the soil’s role in provisioning many essential ecosystem services, including nutrient cycling, water filtration, and sustaining plant productivity (Smith et al., 2015), understanding the complex mechanisms that influence soil C gains and losses is crucial to global change mitigation efforts. This includes examining processes at a variety of spatial and temporal scales, from interactions between soil microbes at the molecular level, who are important mediators of C storage in soil organic matter (SOM), to large-scale ecosystem dynamics that occur over decades or centuries.

Conceptualizing SOM into physically and functionally distinct fractions, including particulate (POM) and mineral-associated organic matter (MAOM), is helpful in understanding how soils might respond to global change and the mechanisms that can be leveraged to enhance the capacity of the soil to store C (Cotrufo and Lavallee, 2022). POM is made of polymeric compounds derived primarily from fragmentation of structural plant inputs, whereas MAOM is formed via sorption of low molecular weight (LMW) soluble compounds (i.e., dissolved organic matter; DOM) to soil mineral surfaces (Yu et al., 2022). While the structural compounds composing POM require enzymatic depolymerization before they can be taken up by microbes, the LMW compounds that serve as precursors to MAOM can be readily taken up and metabolized by microbes if found in solution. However, when bound to soil minerals in MAOM, they are strongly protected from microbial degradation, giving MAOM a longer average residence time than POM (Lavallee et al., 2020; Heckman et al., 2023). Soil microbes play a key role in mediating the formation and persistence of both these fractions. Structural components of microbial necromass such as cell walls can serve as precursors to POM (Cotrufo et al., 2022), while the soluble components of cells and compounds derived from POM depolymerization can sorb to mineral surfaces to form MAOM (Kallenbach et al., 2016; Liang et al., 2019). In fact, microbial necromass may account for as much as half of the total MAOM pool in some ecosystems (Angst et al., 2021; Whalen et al., 2022).

When it is not protected by aggregation, POM is especially susceptible to decomposition via enzyme activity, whereby conditions that support microbial decomposition, including higher temperatures and optimal pH, lead to lower POM storage than in soils in colder climates or acidic soils (Hansen et al., 2024). As such, in both life and death, the structural and functional characteristics of microbes mediate C accumulation and loss in POM and MAOM.

Requiring the availability of active mineral surfaces to form, MAOM is additionally controlled by saturation dynamics, whereby the accumulation of new MAOM is limited by the proportion of mineral surfaces available for organic matter sorption (Hassink, 1997; Six et al., 2002, 2024; Stewart et al., 2007, 2008; Cotrufo et al., 2019; Georgiou et al., 2022; Georgiou et al., 2025). Though exact saturation limits are debated (Begill et al., 2023; Cotrufo et al., 2023; Salonen et al., 2023), many modeling and experimental studies concur that soils that are low in MAOM relative to the available active mineral surface area (i.e., are undersaturated) tend to accumulate more new C in response to inputs than those that are closer to saturation limits (Stewart et al., 2007, 2008; Georgiou et al., 2022). Patterns of C accumulation in undersaturated soils have been fairly robustly explored, and are influenced by a combination of soil properties, C inputs, and management practices. Specifically, besides texture (Hassink, 1997; Cotrufo et al., 2019), mineralogy (Georgiou et al., 2022; King et al., 2023), net primary production (Poeplau et al., 2024), and management (West and Six, 2007) also play a role.

In comparison to soil properties, plant inputs, and management, relatively little work has investigated the extent to which microbial function influences patterns of C accumulation in undersaturated soils. Microbial functions related to how microbes access and metabolize C substrates may be especially relevant in these soils, whereby soils with high matrix capacity force microbes to compete with minerals for access to DOM. Evidence that sorption of inorganic nutrients to mineral surfaces limits their uptake by microbes (Zhu et al., 2016) suggests that when soils are unsaturated, meaning that the majority of mineral sorption sites are unoccupied and are therefore available for DOM sorption, DOM may become limiting to microbes. In turn, microbial communities may require to more enzymes in order to depolymerize structural substrates (e.g., POM) to meet their metabolic needs. This could mean that microbial functions, particularly those related to enzyme production, in soils with low DOM availability (e.g., due to high sorption of soluble substrates driven by undersaturation) compared to soils where DOM availability is high (e.g., due to high saturation of mineral surfaces) could be a key mediator of whether POM is depolymerized or retained in the soil. Given that many conventionally -managed cropland soils and subsoils are typically undersaturated (Georgiou et al., 2022; 2025), microbial traits related to enzyme production may be particularly important to consider in cropland settings and other sites where management may have led to soil degradation.

This proposed relationship between microbial enzyme activity and POM retention is supported by individual-based modeling work that demonstrates how microbial social dynamics promote retention of structural forms of litter and SOM (Allison, 2005; Kaiser et al., 2015). While the term ‘social dynamics’ broadly refers to situations where individual and collect interests conflictive (e.g., Axelrod and Hamilton, 1981; Crespi, 2001; West et al., 2007; Cremer et al., 2019), we define them in the present study specifically as exploitative dynamics between microbes that produce enzymes at maximal capacity (i.e., “producers”) and those that do not produce any enzymes at all (i.e., “cheaters,” who are thus dependent on the activity of enzymes synthesized by producers and exploit them for survival). In Kaiser et al., hypothetical microbial communities comprised entirely of producers had high rates of enzyme production and turnover, regardless of the amount of soluble resources that were available for immediate uptake. This inefficient use of DOM created faster breakdown of structural SOM, and therefore low retention of that SOM pool. On the other hand, microbial communities with cheaters gained the ability to downregulate enzyme production at the community level, due to a feedback mechanism driven by exploitative interactions between the cheaters and enzyme producers. This community-driven feedback mechanism works like a self-regulating control loop: if enzymatic activity generates DOM in excess, it goes to the benefit of cheaters, who do not need to pay the costs for enzyme production. This in turn increases the proportion of cheaters within the community, thereby lowering the total amount of enzymes produced at the community level. If DOM becomes limiting, however, the proportion of cheaters will decrease. Due to the ongoing adjustment of the ratio between enzyme producers and cheaters, the system eventually downregulates its overall enzyme production rate to the minimum necessary to sustain the community (Kaiser et al, 2015).

Compared to hypothetical communities with only producers, those that also contained cheaters produced more necromass that could be recycled within the system relative to the DOM produced by enzymatic breakdown that could be lost via leaching. As demonstrated in Kaiser et al. 2015, the presence of cheaters led to more efficient use of available resources with less waste, conferring to overall higher community carbon use efficiency (CUE), and ultimately slowing decomposition of structural SOM and lowering overall loss of C from the system. Additionally, by increasing the amount of N-rich substrates (i.e., necromass) in the system, cheater presence lowered the overall C:N of DOM produced from the decomposition of both the N-rich necromass and the initial structural substrate that was relatively high in C:N. Thus, the presence of cheaters also allowed the microbial community to overcome any N limitation that may occur due to high initial C:N (Kaiser et al., 2014, 2015). This importance of enzyme regulation to both emergent community metrics like CUE and the fate of different C pools is broadly echoed in other modeling studies. For instance, trade-offs between enzyme production and CUE (Calabrese et al., 2022), feedbacks between substrate availability and enzymes (Sihi et al., 2016), and interactions among enzyme production and diffusion of substrates (Allison, 2005; Abs et al., 2020) all have impacts on soil C decomposition, microbial interactions, and community diversity, highlighting their broad ecological significance (Folse & Allison, 2005; Guseva et al., 2024). Ultimately, given that undersaturation of MAOM may also stress microbial communities, particularly with regards to DOM availability and necessity to produce enzymes, the above-described interactions between community-level enzyme production, structural substrate C:N, and DOM indicate that the relative abundance of producers versus cheaters may play a role in dictating how long structural SOM forms (i.e., POM) are retained in soils that vary in degree of MAOM saturation (i.e., DOM availability), in addition to variations in POM C:N.

Despite experimental evidence of microbial necromass contributions to C storage in POM and MAOM (Kallenbach et al., 2016; Liang et al., 2019; Haddix et al., 2020), as well as frameworks of microbial functional contributions to C storage (e.g., Malik et al., 2020), our empirical understanding of the mechanisms driving POM and MAOM formation and persistence lacks detailed insights into microbial functionality, including emergent behaviors resulting from interactions between microbes that differ in enzyme production capacity. Studies that do not incorporate microbial or enzyme traits explain low variability in POM and MAOM carbon stocks on a global scale (Hansen et al., 2024), suggesting that these traits might account for some of the unexplained variation in soil C storage. More knowledge about how specific microbial interactions, including exploitative interactions between enzyme producers and cheaters, influence C accumulation and loss in the soil would enhance our mechanistic understanding of SOM fraction storage, with broader impacts that could ultimately enable the development of robust, microbe-centric land management strategies aimed at maximizing the retention of C inputs into the soil.

To begin filling this knowledge gap, we used the Kaiser et al. (2015) individual-based model that simulates emergent behaviors of interactions between microbial enzyme producers and cheaters to investigate the extent to which these interactions influence POM decomposition under varying degrees of MAOM saturation. *Sensu* Kaiser et al. (2015), we hypothesized that communities containing both enzyme producers and cheaters would exhibit greater POM retention compared to communities consisting of producers only, where exploitative interactions between producers and cheaters do not exist. Additionally, for communities that contain cheaters, we hypothesized that in saturated soils, high availability of immediately assimilable DOM will require less enzyme investment, leading to enhanced community CUE as well as greater POM retention. By contrast, in undersaturated soils with low DOM availability, we expected that higher enzyme production (and thus a higher proportion of producer biomass) is needed to maintain the microbial community, and POM will be decomposed faster compared to when soils are saturated. Finally, we hypothesized that the C:N of POM would modulate responses to MAOM saturation, with higher POM C:N requiring greater microbial enzyme investment to meet cell stoichiometric needs than when POM C:N is low.

## 2. Materials and Methods

### 2.1. Individual-based modeling

We applied a previously developed, individual-based, and spatially-explicit microscale model (Figure 1; Kaiser et al., 2015) to understand how microbial social dynamics influence emergent community behavior and POM retention under varying degrees of MAOM saturation. Individual-based models simulate the behavior and interactions of individual entities or ‘agents’ within a system. The main objective of these models is to observe how microscale interactions between individual agents give rise to larger scale, emergent properties and behaviors. Our model simulates a 1 x 1 mm^2^ piece of decomposing organic matter as a grid of 100 x 100 microsites or agents. Each microsite measures 10 x 10 x 10 μm and contains a microbial growth model that simulates decomposition of different types of organic matter (Figure 1). These include complex, plant-derived substrate (i.e., plant-derived POM) and microbial-derived C-rich and N-rich substrates, which represent the remains of microbes after cell death (i.e., microbial-derived POM or necromass). Soluble components of dead microbial cells that do not require enzymes to be metabolized contribute directly to the model’s DOM pool following microbial cell death. While plant- and microbial-derived POM pools have a fixed C:N ratio, DOM C:N is dynamic throughout the simulation and varies with the accumulation and degradation of POM (Table 1). At model initialization, 98.5% of the initial total POM pool consists of plant-derived POM and 1.5% consists of microbial necromass. There is also a pool of dissolved inorganic nitrogen (DIN) that can change in size depending on microbial immobilization and mineralization rates, two processes that respond to stoichiometric imbalances between C and N uptake. There is no new input of plant-derived POM throughout the simulations, though microbial-derived POM can accumulate depending on microbial mortality and necromass decomposition rates. Model simulations end when the total amount of substrate is too low or too spatially distant to support microbial activity, and all microbes die.

**Figure 1.**
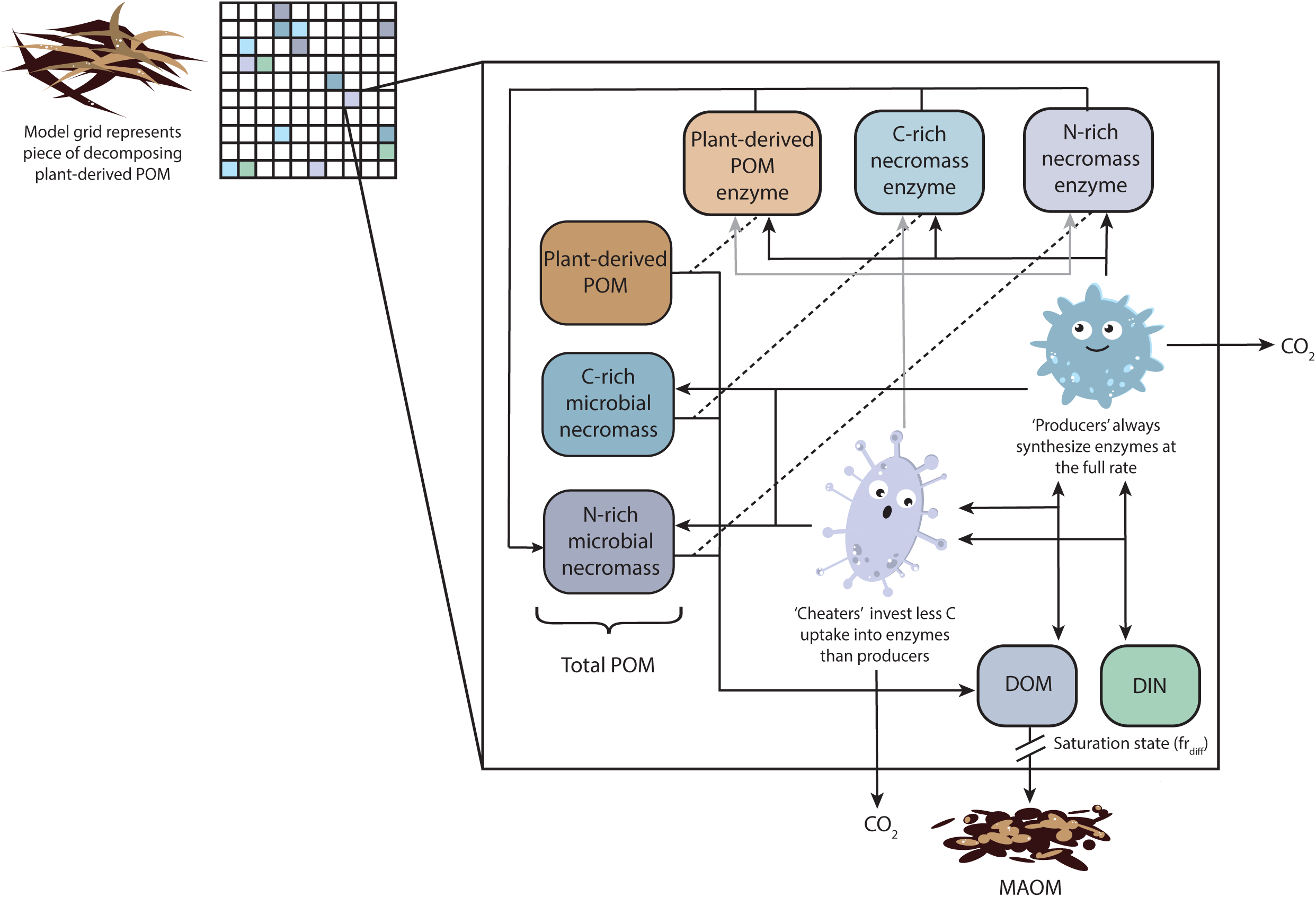

**Table 1.** Key parameters controlling microbial physiology and substrate availability, reproduced following Kaiser et al., 2015, with modifications to match the current model. Additional information about model structure, assumptions, and equations can be found in Table S1, Kaiser et al., 2015, and the supplementary materials therewithin.

| Parameter | Description | Value |
| --- | --- | --- |
| *Microbial cell composition and stoichiometry* |  |  |
| $MB_{fDOM}$ | Fraction of cell biomass accounting for soluble substrates available for immediate uptake by living microbes upon cell death (C:N=15) | 0.06 |
| $MB_{fCR}$ | Fraction of cell biomass accounting for C-rich, complex necromass (e.g., cell wall compounds, lipids, starch; C:N=150) | 0.57 |
| $MB_{fNR}$ | Fraction of cell biomass accounting for N-rich, complex necromass (e.g., proteins, DNA, RNA; C:N=5) | 0.37 |
| $CN_{MB}$ | Resulting microbial biomass C:N | 12.22 |
| *Microbial physiology and enzyme production* |  |  |
| $MB_{fE}$ | Fraction of C uptake invested into enzyme production, following deduction for maintenance respiration | |
|  | Producers | 0.12 |
|  | Cheaters | 0.0 |
| $E_{fPOM}:E_{fCR}:E_{fNR}$ | Ratio in which enzymes are produced for degradation of plant-derived POM:C-rich microbial necromass:N-rich microbial necromass | |
|  | Producers | 0.7:0.15:0.15 |
| *Microbial cell size and turnover* |  |  |
| $MB_{Smax}$ | Maximum microbial cell size, at which a cell divides and colonizes a neighboring microsite | 100 fmol C |
| $MB_{Smin}$ | Minimum microbial cell size, at which a cell dies from starvation | 10 fmol C |
| $MB_{nC}$ | Maximum number of microbial cells per microsite | 1 |
| *Initial pool size and C:N ratio in each microsite* |  |  |
| $C_{POM\_plant}$ | Initial size of the plant-derived POM pool in each microsite | 8333 fmol C (i.e., 83.33 nmol C across entire grid) |
| $CN_{POM\_plant}$ | Initial C:N of plant-derived POM | 10-100, depending on scenario* |
| $C_{POM\_CR}$ | Initial size of the C-rich microbial-derived POM pool in each microsite | 100 fmol C (i.e., 1 nmol C across the entire grid) |
| $CN_{POM\_CR}$ | Initial C:N of C-rich microbial-derived POM (e.g., cell wall compounds, lipids, starch) released upon microbial cell death | 150 |
| $C_{POM\_NR}$ | Initial size of the N-rich microbial-derived POM pool in each microsite | 30 fmol C (i.e., 0.3 nmol C across the entire grid) |
| $CN_{POM\_NR}$ | Initial C:N of N-rich microbial-derived POM (e.g., proteins, DNA, RNA) released upon microbial cell death | 5 |
| $C_{DOM}$ | Initial size of the DOM pool in each microsite | 7 fmol C (i.e., 0.07 nmol C across the |
|  |  | entire grid) |
| $CN_{DOM}^{**}$ | Initial C:N of DOM<sup>**</sup> | 8 |
| $CN_{N-DOM}$ | C:N of soluble necromass released upon microbial cell death | 15 |
| *MAOM formation rates* |  |  |
| $fr_{diff}^*$ | Parameter controlling the fraction of diffusing DOM at each timestep that is capable of sorbing to mineral surfaces; input to Eq. 2 and determined by $fr_L$ (Eq. 3) | 0.0087-0.058, depending on scenario<sup>*</sup> |

<sup>\*</sup>Parameter values altered from Kaiser et al., 2015.
<sup>\*\*</sup> $CN_{DOM}$ is a dynamic parameter varying throughout model simulations.

Depending on functional group (see section 2.2, *Model scenarios*), microbes synthesize three separate enzymes that degrade one of the three structural substrates within our model (i.e., plant-derived POM, C-rich, and N-rich necromass) following Michealis-Menten kinetics:

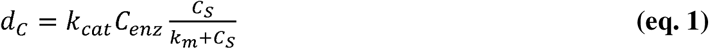

Where *d_C_* is the amount of C released by the enzyme-catalyzed reaction in the microsite within one timestep (i.e., 1 h), *k_cat_* is the number of reactions catalyzed per enzyme per timestep, *C_S_* is the amount of substrate within the microsite, *C_enz_* is the amount of enzyme within the microsite, and *k_m_* is the half saturation constant for an enzyme on its given substrate (Tables 1, S1; Figure 1; Kaiser et al., 2015).

Decomposition products from these reactions, along with the soluble compounds released following microbial cell death, contribute to the DOM pool and are thus available for immediate uptake (Table 1; Figure 1). All enzymes decay following a first order rate constant of 0.036 timestep^-1^ and contribute to the N-rich microbial necromass pool after inactivation (Figure 1).

At model initialization, microbes begin at half of their maximum cell size and are randomly distributed across the grid. Throughout the simulation, each microbial cell takes up DOM and DIN according to their availability within local microsites as well as cell-specific maximum uptakes rates. After uptake, microbes must first meet maintenance respiration needs (Table S1). After maintenance respiration, a functional group-specific proportion of the remaining C uptake is invested into enzyme production (Table 1; see section 2.2, *Model scenarios*). Any C and N remaining after both maintenance and enzyme production is invested into growth. Microbial cells that grow larger than their maximum size reproduce and can colonize empty neighboring microsites (Table 1) or invade an already-occupied microsite (Table S1). Conversely, microbes die when their biomass reaches a lower biomass limit (*MB_Smin_*; Table 1), after failing to access substrate and are thus required to perform maintenance metabolism from their own existing biomass. After death, their necromass is released into the C- and N-rich microbial-derived POM pools, as well as the DOM pool. In addition to starvation, microbes experience random catastrophic death through a stochastic mortality rate (Table S1). After maintenance, enzyme production, and growth needs are met, any stoichiometric imbalances between the amount of C and N acquired are accounted for by either overflow respiration (i.e., in the case of excess C), or by N mineralization or immobilization (i.e., excess N released into or taken up from the DIN pool; Figure 1).

At each time step, 8/9 of the total DOM and DIN in each microsite diffuse to its eight neighboring microsites such that 1/9 remains in the original microsite. A fraction of the total amount of diffusing DOM is lost from the system entirely through sorption to mineral surfaces (i.e., MAOM formation; Figure 1), following:

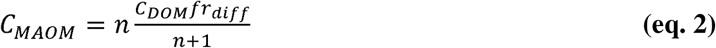

Where *C_MAOM_* is the amount of C leaving the system via MAOM formation, *n* is the number of neighboring microsites (i.e., *n*=8), *C_DOM_* is the total amount of DOM in the microsite, and *fr_diff_* is the fraction of diffusing DOM per timestep (Kaiser et al., 2015). Values of *fr_diff_* are determined following:

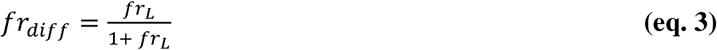

Where *fr_L_* is analogous to the same parameter as described in Kaiser et al., 2015, where it was used to represent leaching from the grid. However, given that leaching and sorption of DOM to mineral surfaces both represent loss of available DOM to microbes within the system, we use *fr_diff_*here to describe saturation state, or the potential of the system to form new MAOM (Figure 1). Note that sorption of DOM to minerals depends on both the free mineral surface area (indicating saturation state) and the amount of diffusing DOM in the environment. In this context, *fr_diff_* acts as a relative parameter, where the amount of MAOM formed at each timestep depends not just on the value of *fr_diff_*, but also on the amount of DOM that is diffusing across the grid and how much new DOM is produced by the activity of any enzymes present in the system. We varied *fr_diff_* to simulate the effects of differences in the potential to form new MAOM on microbes, where low values of *fr_diff_* approximate low potential for sorption of DOM to mineral surfaces when soils are already saturated with MAOM, and high values of *fr_diff_* approximate high potential of DOM sorption to minerals when soils are unsaturated (see section 2.2, *Model scenarios*). Importantly, we assume that the amount of DOM sorbing to minerals does not significantly alter the state of MAOM saturation during our simulations, and therefore *fr_diff_*is kept constant throughout each simulation.

Additional information about model structure, assumptions, and equations can be found in Table S1, Kaiser et al., 2015, and the supplementary materials therewithin.

### 2.2. Model scenarios

We compare model scenarios in which all microbes produce extracellular enzymes at the full rate (i.e., 0.12 of the C uptake at each timestep is invested into enzyme production, after deductions for maintenance respiration) with scenarios in which only half of the microbial community produces enzymes. We define the microbes that do not produce any enzymes as “cheaters,” and those that do as “producers” (Table 1). Given that cheaters do not produce any enzymes, their survival relies to at least some extent upon access to DOM that has already been produced by the enzyme activity of producers.

Outside of enzyme production, all microbes invest the same amount of acquired resources into cellular maintenance and growth and possess identical cell chemical composition and stoichiometry (Table 1; Figure 1). Note that while trade-offs between enzyme production and growth, including return-on-investment for new resource acquisition, are key aspects of microbial physiology, they have been robustly explored in other modeling studies (e.g.,Allison, 2005; Calabrese et al., 2022) and are therefore not a focus of this study. As such, our work considers a direct comparison of communities that contain microbial cheaters *versus* those that contain only producers, without introducing any other axes of variation in microbial growth, resource acquisition, or life history traits.

In addition to enzyme production, we compare scenarios varying the C:N ratio of plant-derived POM inputs as well as the rate of new MAOM formation. We consider plant-derived POM C:N ratios from 10-100, in intervals of 10, corresponding to different input qualities spanning from the low C:N of leguminous plants to the high C:N of woody plants. We also consider seven different MAOM saturation states by altering the fraction of soluble C leaving the system (*fr_diff_*) through consideration of six different values of *fr_L_* (i.e., 0.0088, 0.0176, 0.0264, 0.0352, 0.044, 0.0528, and 0.0616; Eq. 3; Table 1), corresponding to *fr_diff_* values of 0.0087, 0.0173, 0.0257, 0.034, 0.0421, 0.0502, and 0.0508 (Table 1; Figure 1). We interpret high potential for MAOM formation (i.e., loss of diffusible C) to be representative of an undersaturated soil, where a higher proportion of diffusing DOM is inaccessible to microbes, while low potential for MAOM formation represents a soil that is already saturated with respect to MAOM, where a higher proportion of DOM is available for immediate uptake.

For both communities that did and did not contain cheaters, we ran simulations in fully factorial combinations of the initial plant-derived POM C:N and MAOM saturation (i.e., potential to form new MAOM) parameter values described above. To account for the stochastic nature of the model, we ran ten replicate simulations of each initial plant-derived POM C:N and MAOM saturation combination for simulations containing only enzyme producers and for simulations containing both producers and cheaters. However, some of the above combinations of plant-derived POM C:N and MAOM saturation were strongly limiting to microbes, causing the full community to die before they were able to begin degrading the initial POM input. This was particularly true for model scenarios of both producers and cheaters when POM C:N and *fr_diff_*were greater than 80 and 0.0502, respectively. We excluded any such replicates from downstream analyses. Further details on how many replicates were excluded from specific combinations of POM C:N and *fr_diff_* are located in Table S2.

### 2.3. Model outputs

We investigated the effects of interactions between microbial enzymes producers and cheaters on hypothetical soils that vary in both initial, plant-derived POM C:N and MAOM saturation by evaluating how sensitive cumulative MAOM formation was to variations in both initial POM C:N and *fr_diff_*(i.e., saturation state). We also assessed how multiple community properties responded to initial POM C:N and *fr_diff_*, including total enzyme production, total microbial biomass, the ratio of enzyme production to biomass (i.e., enzymes:total biomass; indicative of proportional enzyme investment needed to support the community), community CUE (Manzoni et al., 2012; Kaiser et al., 2015), net growth rate, and for simulations that contained cheaters, the ratio of cheater to total microbial biomass (i.e., cheater:total biomass). Following Kaiser et al., each of these microbial metrics were calculated from C fluxes aggregated across the grid, such that each represents an emergent property of the full microbial and soil system. In particular, community CUE was calculated following:

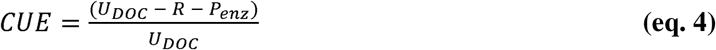

Where *U_DOC_* is the total amount of dissolved organic carbon (DOC) taken up by all of the microbes on the grid, *R* is the total amount of C respired by all microbes, and *P_enz_* is the total amount of C released as enzymes (Manzoni et al., 2012; Kaiser et al., 2015). Because the model’s behavior is highly random when microbes become limited by resource availability, especially towards the end of the simulation, we considered values of these variables when 60% of the total initial C pool was degraded (Kaiser et al., 2015). Lastly, to understand how interactions between producers and cheaters, as well as variation in both plant-derived POM C:N and MAOM saturation affect decomposition rates of the total POM pool (i.e., both plant- and microbial-derived POM), we assessed how the amount of time it took the community to decompose 60% of the initial C pool was affected by both initial POM C:N and *fr_diff_*.

## 3. Results

### 3.1. Sensitivity of cumulative MAOM formation to cheater presence, initial plant-derived POM C:N, and frdiff

We first sought to understand whether relationships between *fr_diff_* (i.e., MAOM saturation state), initial plant-derived POM C:N, and the cumulative amount of MAOM formed differed between simulations that contained only enzyme producers, and those that also contained cheaters. Model scenarios that contained only producers exhibited greater cumulative MAOM formation than those that contained both producers and cheaters, and largely responded only to changes in initial plant-derived POM C:N (Figure 2). Specifically, cumulative MAOM formation in producer-only scenarios decreased with increases in POM C:N, particularly when C:N was greater than 30. On the other hand, cumulative MAOM formation in simulations that contained both cheaters and producers was sensitive to both *fr_diff_* and initial, plant-derived POM C:N. In these scenarios, cumulative MAOM formed increased with increasing values of *fr_diff_* and decreased slightly with increases in POM C:N, especially when it was greater than 30 (Figure 2). Additionally, within individual levels of *fr_diff_*, lower initial, plant-derived POM C:N values led to greater cumulative MAOM formation than when initial C:N was high (Figure 2).

**Figure 2.**
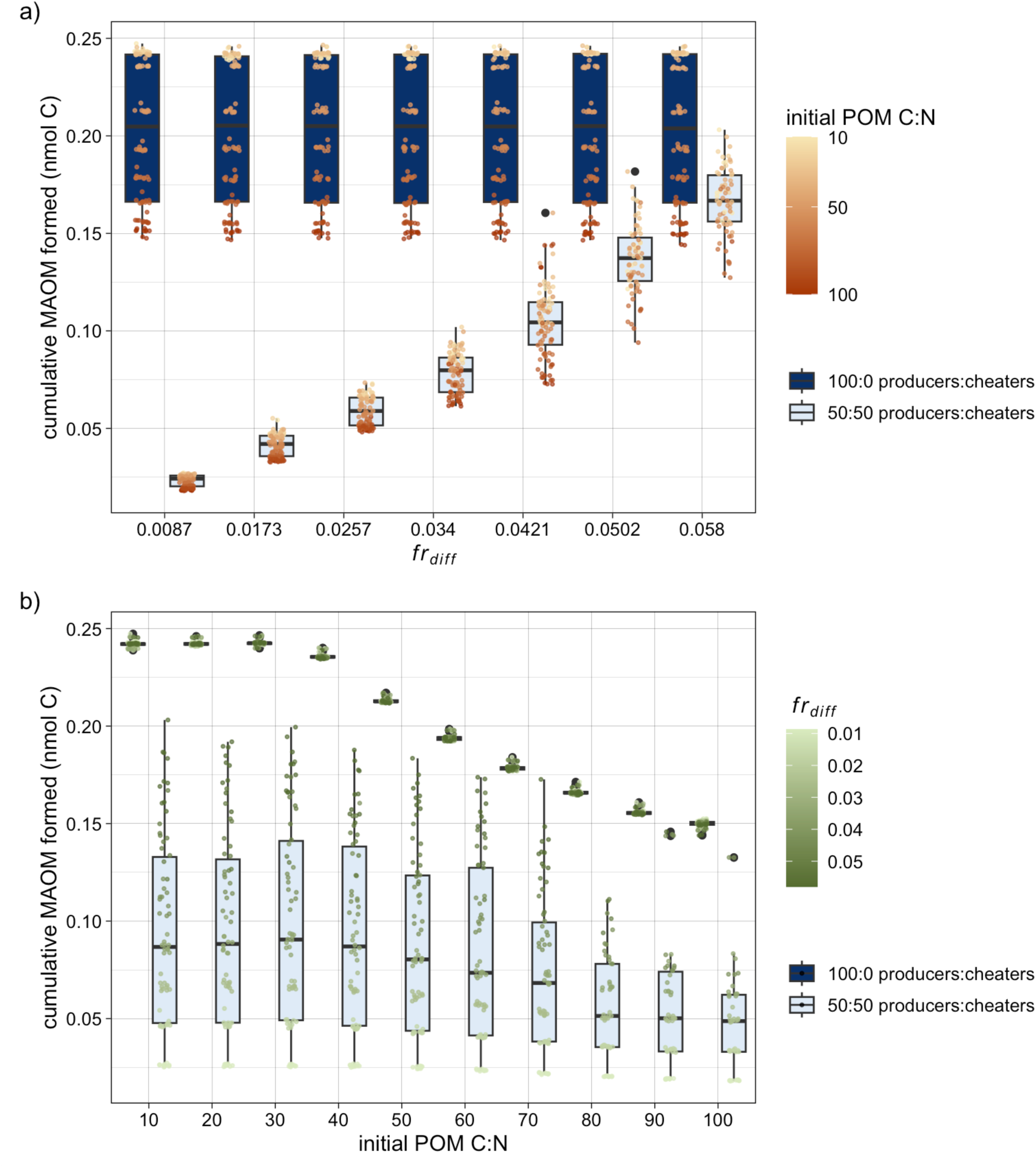

### 3.2 Microbial community responses to initial plant-derived POM C:N and fr_diff_

Emergent microbial community responses to initial plant-derived POM C:N and *fr_diff_* (i.e., MAOM saturation state) depended on whether simulations consisted of only enzyme producers, or contained both producers and cheaters. In general, emergent community properties in simulations that contained only producers responded solely to changes in plant-derived POM C:N, with increases in C:N leading to greater enzyme production relative to total biomass (i.e., enzymes:total biomass), but decreased community CUE, net growth rate, total biomass, and total enzyme production (Figures 3-4, Figure S1).

**Figure 3.**
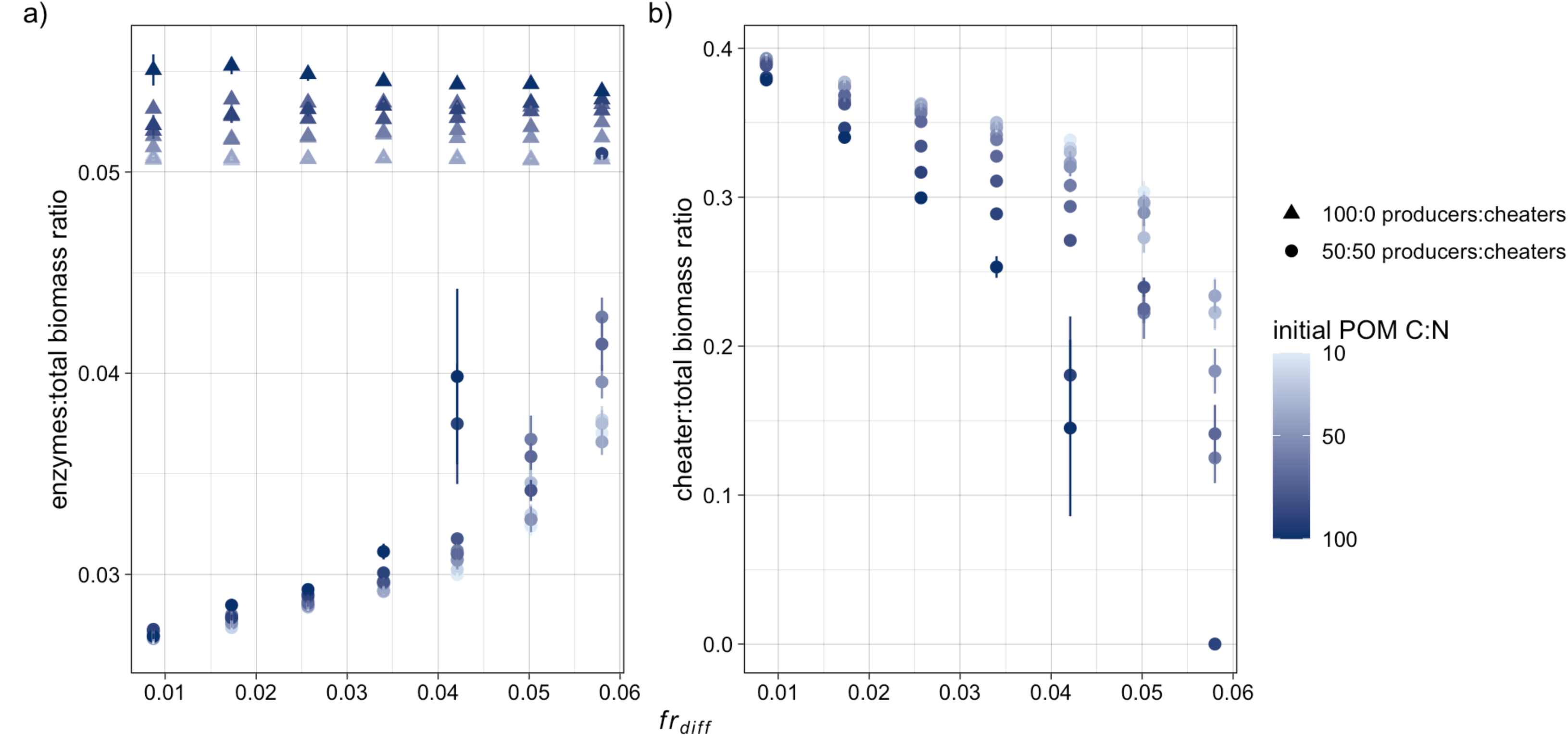

**Figure 4.**
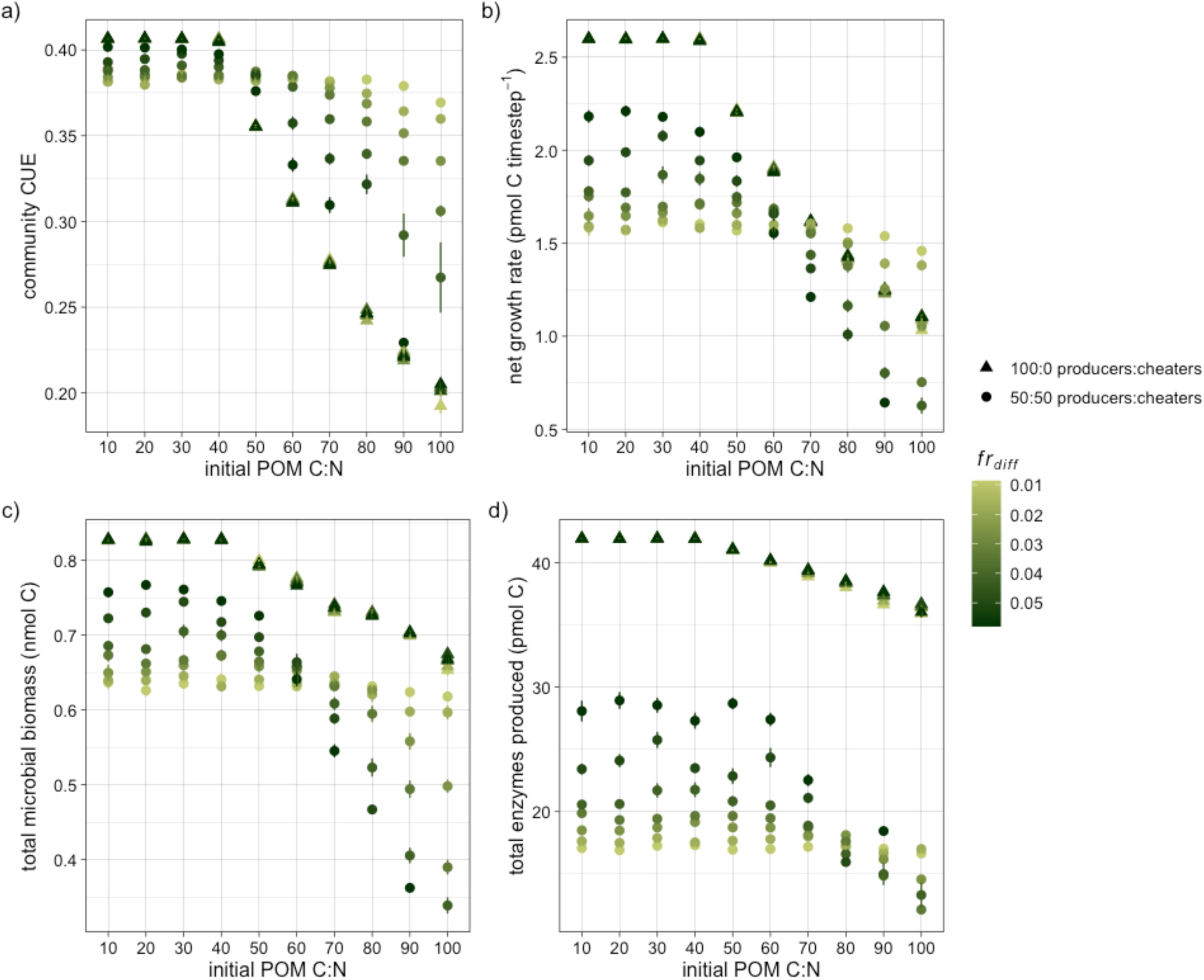

In contrast, emergent community properties in simulations that contained cheaters and enzyme producers were responsive to changes in both initial plant-derived POM C:N and *fr_diff_*. Proportional enzyme investment (i.e., enzymes:total biomass ratio) in scenarios containing cheaters was lower than that of producer-only scenarios, and generally increased in response to *fr_diff_*. However, the rate at which enzymes:total biomass increased with *fr_diff_* depended on POM C:N, with higher values of initial plant-derived POM C:N conferring to steeper increases in enzymes:total biomass than under low initial C:N ratios (Figure 3a, Figure S1). Similarly, while cheater:total biomass generally declined with increasing values of *fr_diff_*, the rate at which cheater biomass decreased depended on plant-derived POM C:N. Higher C:N ratios led to steeper declines in cheater:total biomass in response to *fr_diff_* than when C:N was low (Figure 3b, Figure S1).

While community CUE, net growth rate, total microbial biomass, and total enzymes produced in simulations that contained producers and cheaters also depended on both initial plant-derived POM C:N and *fr_diff_*, the dynamics of these responses differed markedly from those of proportional enzyme investment and cheater biomass (Figure 4; Figure S2). Generally, increases in plant-derived POM C:N were associated with declines in CUE, growth, total biomass, and enzymes. However, when plant-derived POM had a relatively low C:N, increases in *fr_diff_* had an overall positive effect on each of the above variables, leading CUE, growth, biomass, and enzyme production. This relationship reversed when POM C:N was high, such that increases in *fr_diff_* had an overall negative effect. While the presence of this reversal was consistent across CUE, growth, total biomass, and enzyme production, the specific C:N where the reversal occurred depended on microbial metric. For instance, reversal of the impacts of *fr_diff_*occurred around a C:N of 50 for community CUE, but around a C:N of 80 for total enzyme production (Figure 4; Figure S2).

### 3.3. Effects of initial plant-derived POM C:N and fr_diff_ on the decomposition of the total initial C pool

Regardless of initial plant-derived POM C:N and *fr_diff_*, communities that contained only enzyme producers decomposed 60% of the total initial C pool two to four times faster than communities that consisted of both producers and cheaters (Figure 5; Figure S3). The amount of time it took for producer-only communities to degrade 60% of the initial C pool depended solely on plant-derived POM C:N, with increases in C:N leading to slightly slower decomposition of the initial substrate. When communities also contained cheaters, decomposition rates depended on both POM C:N and *fr_diff_*. For these simulations, increases in plant-derived POM C:N generally led to slower decomposition of the initial C pool.

**Figure 5.**
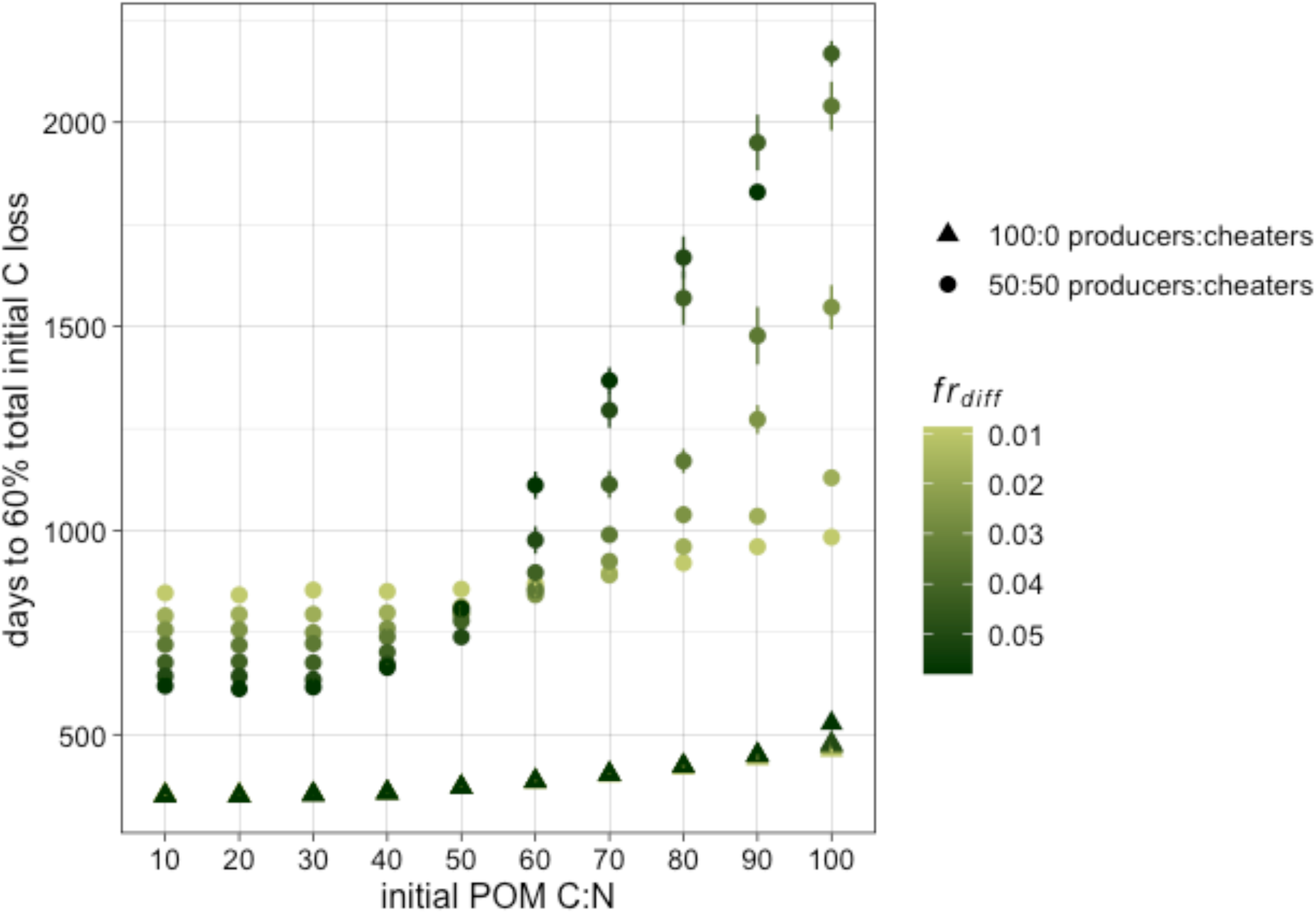

However, *fr_diff_* and plant-derived POM C:N interacted with one another such that *fr_diff_* had a negative effect on the amount of time it took the community to degrade 60% of the initial C input when POM C:N was less than ∼60, but a positive effect when C:N was greater than 60. Additionally, decomposition responded more strongly to changes in plant-derived POM C:N when *fr_diff_* was high, with decomposition rates with respect to POM C:N decreasing much more rapidly under high values of *fr_diff_* compared to low (Figure 5; Figure S3).

## 4. Discussion

In this study, we used individual-based modeling to understand the potential for interactions between microbes that do and do not produce enzymes to influence POM decomposition and new MAOM formation in hypothetical soils that vary in their degree of MAOM saturation, simulated by altering the potential of the model system to form new MAOM (i.e., *fr_diff_*). Consistent with previous findings, we demonstrate that the presence of microbial cheaters led to slower decomposition of structural SOM (i.e., POM) compared to communities that consisted of only enzyme producers (Kaiser et al., 2015).

In addition, we found that our model system responded to changes in MAOM saturation and initial plant-derived POM C:N in unique ways, depending on whether communities contained cheaters. While the system’s dynamics were generally limited only by N availability when it contained just enzyme producers, the limitations of MAOM undersaturation (i.e., high *fr_diff_*) in simulations that contained cheaters altered the capacity of the microbial cheaters to buffer the system against N limitation, with consequences on emergent microbial behaviors and structural C decomposition. Altogether, these findings indicate that microbial traits, especially those related to regulation and production of enzymes, affect not only structural substrate decomposition but also new MAOM formation, and are essential to consider in other empirical and modeling work.

Our results revealed that new MAOM formation is influenced not only by its saturation state, but also by initial POM C:N and, most critically, interplays between microbes that do and do not produce extracellular enzymes. In simulations containing only enzyme producers, where enzyme production was not regulated at the community level (Kaiser et al., 2015), we observed no effect of *fr_diff_*within the analyzed range (0.87 - 5.8%) on total microbial biomass (Fig. 4c), total enzyme production (Fig. 4d), or POM degradation time (Fig. 5). An effect of *fr_diff_*emerged at higher values (e.g., above 7%; results not shown), but at these levels, microbial communities faced a high risk of death due to insufficient DOM availability and often failed to survive beyond ∼30 days. The resilience of producer communities as well as the null effects of *fr_diff_* on cumulative MAOM formation within the analyzed range of this parameter reflect the continued excess production of DOM (Figure S4) due to high, unregulated enzyme production. Additionally, competition for DOM between minerals and microbes in these producer-only simulations was relatively weak. This was because the microbes have preferential access to the products generated by their own enzymes due to their spatial proximity – they can take up the DOM that is produced in their own microsites before it diffuses away and sorbs to minerals. These competitive dynamics between producer microbes and minerals are generally echoed in other individual-based modeling work on the trade-offs faced by producers of extracellular enzymes (Allison, 2005; Guseva et al., 2024). Such studies demonstrate that low enzyme diffusion rates allow the microbes that produced those enzymes to have preferential access to decomposition products, ultimately favoring their growth (Allison, 2005). This may indicate that excess enzyme production, even if it results in more DOM than needed for microbial growth, could give enzyme producers an advantage over their competitors for DOM, regardless of whether those competitors are other microbes or mineral surfaces available for sorption.

In contrast to simulations that contained only enzyme producers, cumulative MAOM formation in those that also contained cheaters responded to both MAOM saturation and plant-derived POM C:N. In these scenarios, decreases in both MAOM saturation and plant-derived POM C:N led to greater cumulative MAOM formation. The community’s collective ability to regulate enzyme production due to dynamic feedbacks between producers and cheaters made the system more responsive to shifts in MAOM saturation and associated changes in DOM availability. For instance, the proportion of cheaters in the community declined with increasing *fr_diff_* (i.e., decreased saturation), thereby increasing enzyme production at the community level and eventually outbalancing the loss of DOM. Specifically, high saturation (i.e., low *fr_diff_*) necessitated lower enzyme production to maintain the community (reflected by higher proportional cheater biomass), resulting in less DOM that could contribute to MAOM formation.

The reverse was true when saturation was low: this condition required greater enzyme production to break down POM for community maintenance, which diminished cheater presence and led to increased DOM that could ultimately form new MAOM. Furthermore, plant-derived POM C:N acted as an additional limitation on cumulative MAOM formation, as lower N availability, both within specific *fr_diff_* levels and across the entire range tested, consistently led to less MAOM formation than when N availability was higher. This effect was especially pronounced under unsaturated conditions (i.e., high *fr_diff_*), where decreases in POM C:N produced a comparatively larger increase in MAOM formation than under saturated conditions (i.e., low *fr_diff_*). These findings align with empirical research on interactions between MAOM saturation and N availability (Wu et al., 2022), and lends that MAOM saturation state may be a stronger predictor of new mineral-stabilized C accumulation than the nutrient status of incoming C inputs.

The above dynamics of cumulative MAOM formation are generally mirrored in the responses of emergent microbial properties to initial plant-derived POM C:N and MAOM saturation. In support of our hypotheses and similar to cumulative MAOM formation, we found that saturation and plant-derived POM C:N interacted in complex ways to shape community dynamics when communities contained both cheaters and enzyme producers. Overall, increases in saturation (i.e., lower *fr_diff_*) resulted in higher cheater biomass and declines in proportional enzyme production. In these cases, reduced DOM availability under low saturation made it harder for cheaters to survive, as greater enzyme investment was needed to break down POM for the community to meet its metabolic needs. While this general response was consistent across all tested initial POM C:N ratios, its strength depended on C:N, with higher C:N ratios leading to more rapid decreases in cheaters and corresponding increases in enzymes, in proportion to total biomass, than under low C:N. This indicates that communities consisting of both enzyme producers and cheaters were co-limited by N availability and MAOM saturation. While microbial colimitation has been well-studied in a variety of contexts (e.g., Wutzler and Reichstein, 2008; Choi et al., 2022; Held et al., 2024; Cui et al., 2025), our modeling experiments indicate that the state of MAOM saturation may represent another limiting factor on microbial growth, whereby greater plant-derived POM C:N (or other stressors) enhance the negative effects of MAOM undersaturation on the community.

Despite the above, findings from our analysis of community CUE, net growth rates, total biomass and enzyme production, as well as decomposition rates, indicate that colimitation of N availability and MAOM undersaturation may be more nuanced than indicated with relative enzyme production and relative cheater biomass. For each of the former variables, increases in MAOM formation had an overall positive effect when plant-derived POM C:N was low (e.g., below ∼50), but a negative effect when C:N was high. This indicates our original hypothesis – that undersaturation leads to greater enzyme production due to competition between microbes and mineral surfaces for DOM, and therefore faster decomposition of structural SOM – is only true when the microbial community is not limited by N availability. High MAOM formation rates act to reduce the number of cheaters the community can support, thereby decreasing the flexibility of the system to respond to N limitation via regulation of enzymes and making the system more sensitive to changes in N availability. In other words, while a high capacity to maintain cheaters (e.g., when MAOM saturation is high) has an overall negative effect on the system when N availability is high, it has a positive effect when N is low. At this point, cheaters lose their buffering capacity, and the system begins to more closely resemble those that contain only enzyme producers in that it becomes characterized by steep declines in CUE, growth, total biomass, total enzymes, and decomposition rates in response to increases in C:N.

However, POM in most environments rarely has a C:N above ∼50 (unless it is derived mostly from woody litter; Cotrufo et al., 2019; Lugato et al., 2021; Yu et al., 2022), making it unlikely that microbial cheaters in typical, unsaturated soils lose their buffering capacity solely due to POM-derived N limitations. However, our findings suggest broader implications for how environmental stressors may influence the buffering capacity of cheaters, regardless of which specific environmental factor may be limiting microbial growth. Microbial growth and activity can be constrained by multiple factors simultaneously, including but not limited to soil moisture, temperature, nutrient availability, and accessibility of C substrates (e.g., Elser et al., 1995; Treseder, 2008; Kamble and Bååth, 2016; Castle et al., 2017). In unsaturated soils where microbes are also experiencing additional activity constraints, community dynamics may more closely resemble those of a system composed solely of enzyme producers. Until MAOM storage begins to approach saturation, this may lead to heightened sensitivity of the community to additional stressors, with consequences on the decomposition of structural litter and SOM.

Given the markedly different responses of the system to MAOM saturation and POM C:N depending on cheater presence, our findings underscore the importance of considering microbial traits in studies seeking to understand C changes in soils that differ in MAOM saturation. Our simulated dynamics of simultaneous MAOM formation and POM retention in undersaturated conditions, particularly when microbial communities include cheaters compared to those without, is consistent with findings from several field studies and meta-analyses. For instance, long-term adoption of regenerative agricultural practices, such as no- or low-till, intercropping, livestock integration, and adaptive multi-paddock grazing, in traditionally-managed agricultural soils (e.g., undersaturated soils; Georgiou et al., 2022) has been shown to promote C accumulation in both POM and MAOM (Mosier et al., 2021; Prairie et al., 2023). Similarly, ecosystem restoration efforts, such as converting degraded croplands to native vegetation, have been shown to increase both C fractions concurrently (Kalinina et al., 2019; Yang et al., 2022). In combination with the present study, these findings support the notion that microbial social dynamics and community-level enzyme regulation could be additional drivers of ecosystem-scale patterns of SOM accumulation, in addition to the mechanisms of C accrual identified in previous work (e.g., Hassink, 1997; Stewart et al., 2007, 2008; West and Six, 2007; Cotrufo, 2019; Georgiou et al., 2022).

Although our model was able to approximate patterns of POM and MAOM storage in undersaturated soils, our simulations do not account for several C cycling dynamics that are present in natural systems. These include continuous inputs of new C to the system, as well as microbial decomposition of MAOM (Jilling et al., 2021), a process known to occur particularly under N limitation (Mazzilli et al., 2014). Despite this, high retention of both plant- and microbial-derived POM in our model (i.e., up to ∼6 years to degrade 60% of the total initial input) implies that any structural C entering the system through processes such as litterfall and rhizodeposition may also be retained. Furthermore, the initial plant-derived POM input amount used in our simulations was not designed to cause any microbial limitations by itself, although its variable properties like C:N can (Kaiser et al., 2014, 2015). Lower microbial biomass and a reduced capacity of cheaters to buffer the system indicate that our hypothetical communities were N-limited, especially when both initial plant-derived POM C:N and MAOM formation rates were high. Though our microbes were not able to access any of the MAOM formed throughout the simulation, communities in natural soil systems could meet their N needs by breaking down existing MAOM stores, potentially altering MAOM formation and POM decomposition patterns – particularly when the latter is shaped by the co-limitation dynamics explored in this study. Although exploitative interactions between producers and cheaters can be challenging to empirically quantify, integrating - omics techniques, enzyme assays (e.g., to quantify enzyme production relative to microbial biomass), stable isotope tracing, and other functional measurements (e.g., Malik et al., 2020) into studies on MAOM formation could provide clearer insights into how these limitations affect our analysis. This could also build experimental evidence for the role of interactions between enzymes producers and cheaters in mediating soil C storage, as well as the relative importance of these interactions to C storage in comparison to other aspects of enzyme production, including enzyme energetics and return on investment (Allison, 2005; Calabrese et al., 2022).

Despite these limitations on our analysis, our ability to replicate patterns of simultaneous POM retention and MAOM accumulation observed in field studies underscores the significance of microbial interactions and other community traits that shape enzyme production in driving SOM formation and persistence. These traits and emergent behaviors are critical to consider not just in empirical work, but also in the development of process-based models of soil C and N cycling. While several current models include microbial biomass as a mediator of structural C decomposition (e.g., Abramoff et al., 2022; Chandel et al., 2023; Rocci et al., 2024), relatively few explicitly account for differences in microbial ecology (e.g., Sistla et al., 2014; Wieder et al., 2014; Wang et al., 2015; Georgiou et al., 2017) that are known mediators of C dynamics (e.g., Allison et al., 2010; Trivedi et al., 2013; Kaiser et al., 2014; Crowther et al., 2015; Buchkowski et al., 2017; Hall et al., 2018; Bradford et al., 2021). Our work builds upon calls to better incorporate microbial ecology into ecosystem models (e.g., Rocci et al., 2024), and provides evidence that incorporating microbial interactions and other community traits that can shape both enzyme production and decomposition, such as the interactions between producers and cheaters explored in this study, has the potential to increase process-based model precision and accuracy. This may be especially true in ecosystems where microbial communities may strongly mediate C fraction storage [e.g., mesic environments; (Cotrufo et al., 2021)].

In addition to process-based model development, relationships between microbial enzyme regulation and structural SOM decomposition offer a valuable framework for developing microbe-centric land management strategies that promote C storage and N retention (Kaiser et al., 2015). For example, efforts to develop microbial inoculants for use in agricultural soils may want to focus selection of microbial strains or genes that promote enzyme regulation in response to soluble substrate availability, especially when other environmental stressors are not limiting. In combination with other strategies to increase plant inputs, this approach could maximize microbial community efficiency in ways that retain more C. Alongside other regenerative practices that boost C inputs to the soil, microbial interventions aimed at retaining or efficiently recycling SOM while minimizing waste metabolism may be able to extend the residence time of newly-formed soil C, particularly in the form of POM.

## 5. Conclusions

In this study, we demonstrate that microbial social dynamics, defined as exploitative interactions between microbes that differ in their capacity to produce extracellular enzymes, may be an additional mechanism contributing to observed patterns of POM and MAOM accumulation in undersaturated soils. In particular, emergent microbial properties, POM decomposition, and MAOM formation vary significantly based on the ability of hypothetical microbial communities to regulate enzyme production at the community level, as well as the extent to which MAOM undersaturation, N availability, and other environmental stressors impose co-limitations on the system. While more work is needed to empirically quantify the effects of community-level enzyme regulation on C fraction storage, our study provides a useful roadmap for more comprehensive incorporation of microbial ecology into process-based models of soil C and N cycling, as well as the development of microbial interventions to promote C storage and N retention in managed soils. Ultimately, with an improved understanding of how microbial community functions like enzyme regulation impact SOM fractions, we may be better able to realize the potential of soil microbiomes as a tool to address global change.

## Acknowledgements

We thank members of the Cotrufo Soil Innovation Lab (SoIL) for their helpful suggestions on our analyses and manuscript. This work was supported by a European Research Council-National Science Foundation (ERC-NSF) supplement to NSF DEB award #2016003, as well as by the ERC under the European Union’s Horizon 2020 research and innovation programme (grant agreement No819446 to CK).

## References

Abramoff, R.Z., Guenet, B., Zhang, H., Georgiou, K., Xu, X., Viscarra Rossel, R.A., Yuan, W., Ciais, P., 2022. Improved global-scale predictions of soil carbon stocks with Millennial Version 2. Soil Biology and Biochemistry 164, 108466. doi:10.1016/j.soilbio.2021.108466

Abs, E., Leman, H., Ferrière, R., 2020. A multi-scale eco-evolutionary model of cooperation reveals how microbial adaptation influences soil decomposition. Communications Biology 3, 520. doi:10.1038/s42003-020-01198-4

Allison, S.D., 2005. Cheaters, diffusion and nutrients constrain decomposition by microbial enzymes in spatially structured environments. Ecology Letters 8, 626–635. doi:10.1111/j.1461-0248.2005.00756.x

Allison, S.D., Wallenstein, M.D., Bradford, M.A., 2010. Soil-carbon response to warming dependent on microbial physiology. Nature Geoscience 3, 336–340. doi:10.1038/ngeo846

Angst, G., Mueller, K.E., Nierop, K.G.J., Simpson, M.J., 2021. Plant- or microbial-derived? A review on the molecular composition of stabilized soil organic matter. Soil Biology and Biochemistry 156, 108189. doi:10.1016/j.soilbio.2021.108189

Axelrod, R., Hamilton, W.D., 1981. The Evolution of Cooperation. Science 211, 1390–1396. doi:10.1126/science.7466396

Begill, N., Don, A., Poeplau, C., 2023. No detectable upper limit of minerallJassociated organic carbon in temperate agricultural soils. Global Change Biology gcb.16804. doi:10.1111/gcb.16804

Bradford, M.A., Wood, S.A., Addicott, E.T., Fenichel, E.P., Fields, N., González-Rivero, J., Jevon, F.V., Maynard, D.S., Oldfield, E.E., Polussa, A., Ward, E.B., Wieder, W.R., 2021. Quantifying microbial control of soil organic matter dynamics at macrosystem scales. Biogeochemistry 156, 19–40. doi:10.1007/s10533-021-00789-5

Buchkowski, R.W., Bradford, M.A., Grandy, A.S., Schmitz, O.J., Wieder, W.R., 2017. Applying population and community ecology theory to advance understanding of belowground biogeochemistry. Ecology Letters 20, 231–245. doi:10.1111/ele.12712

Castle, S.C., Sullivan, B.W., Knelman, J., Hood, E., Nemergut, D.R., Schmidt, S.K., Cleveland, C.C., 2017. Nutrient limitation of soil microbial activity during the earliest stages of ecosystem development. Oecologia 185, 513–524. doi:10.1007/s00442-017-3965-6

Chandel, A.K., Jiang, L., Luo, Y., 2023. Microbial Models for Simulating Soil Carbon Dynamics: A Review. Journal of Geophysical Research: Biogeosciences 128, e2023JG007436. doi:10.1029/2023JG007436

Cotrufo, M.F., 2019. Soil carbon storage informed by particulate and mineral-associated organic matter. Nature Geoscience 12, 8.

Cotrufo, M.F., Haddix, M.L., Kroeger, M.E., Stewart, C.E., 2022. The role of plant input physical-chemical properties, and microbial and soil chemical diversity on the formation of particulate and mineral-associated organic matter. Soil Biology and Biochemistry 168, 108648. doi:10.1016/j.soilbio.2022.108648

Cotrufo, M.F., Lavallee, J.M., 2022. Soil organic matter formation, persistence, and functioning: A synthesis of current understanding to inform its conservation and regeneration, in: Advances in Agronomy. Elsevier, pp. 1–66. doi:10.1016/bs.agron.2021.11.002

Cotrufo, M.F., Lavallee, J.M., Six, J., Lugato, E., 2023. The robust concept of mineral-associated organic matter saturation: A letter to Begill et al., 2023. Global Change Biology 29, 5986–5987. doi:10.1111/gcb.16921

Cotrufo, M.F., Lavallee, J.M., Zhang, Y., Hansen, P.M., Paustian, K.H., Schipanski, M., Wallenstein, M.D., 2021. I n lJNlJO ut: A hierarchical framework to understand and predict soil carbon storage and nitrogen recycling. Global Change Biology 27, 4465–4468. doi:10.1111/gcb.15782

Cotrufo, M.F., Ranalli, M.G., Haddix, M.L., Six, J., Lugato, E., 2019. Soil carbon storage informed by particulate and mineral-associated organic matter. Nature Geoscience 12, 989–994. doi:10.1038/s41561-019-0484-6

Cremer, J., Melbinger, A., Wienand, K., Henriquez, T., Jung, H., Frey, E., 2019. Cooperation in Microbial Populations: Theory and Experimental Model Systems. Journal of Molecular Biology 431, 4599–4644. doi:10.1016/j.jmb.2019.09.023

Crespi, B.J., 2001. The evolution of social behavior in microorganisms. Trends in Ecology & Evolution 16, 178–183. doi:10.1016/S0169-5347(01)02115-2

Crowther, Thomas W., Sokol, Noah W., Oldfield, Emily E., Maynard, Daniel S., Thomas, Stephen M., Bradford, Mark A., Crowther, T. W., Sokol, N. W., Oldfield, E. E., Maynard, D. S., Thomas, S. M., Bradford, M. A., 2015. Environmental stress response limits microbial necromass contributions to soil organic carbon. Soil Biology and Biochemistry Complete, 153–161. doi:10.1016/j.soilbio.2015.03.002

Elser, J.J., Stabler, L.B., Hassett, R.P., 1995. Nutrient limitation of bacterial growth and rates of bacterivory in lakes and oceans: a comparative study. Aquatic Microbial Ecology 09, 105–110. doi:10.3354/ame009105

Folse, H.J., Allison, S.D., 2012. Cooperation, Competition, and Coalitions in Enzyme-Producing Microbes: Social Evolution and Nutrient Depolymerization Rates. Frontiers in Microbiology 3. doi:10.3389/fmicb.2012.00338

Georgiou, K., Abramoff, R.Z., Harte, J., Riley, W.J., Torn, M.S., 2017. Microbial community-level regulation explains soil carbon responses to long-term litter manipulations. Nature Communications 8, 1223. doi:10.1038/s41467-017-01116-z

Georgiou, K., Jackson, R.B., Vindušková, O., Abramoff, R.Z., Ahlström, A., Feng, W., Harden, J.W., Pellegrini, A.F.A., Polley, H.W., Soong, J.L., Riley, W.J., Torn, M.S., 2022. Global stocks and capacity of mineral-associated soil organic carbon. Nature Communications 13, 3797. doi:10.1038/s41467-022-31540-9

Georgiou, K., Angers, D., Champiny, R.E., Cotrufo, M.F., Craig, M.E., Doetterl, S., Grandy, A.S., Lavallee, J.M., Lin, Y., Lugato, E., Poeplau, C., Rocci, K.S., Schweizer, S.A., Six, J., Wieder, W.R., 2025. Soil Carbon Saturation: What Do We Really Know? Global Change Biology 31, e70197. doi:10.1111/gcb.70197

Guseva, K., Mohrlok, M., Alteio, L., Schmidt, H., Pollak, S., Kaiser, C., 2024. Bacteria face trade-offs in the decomposition of complex biopolymers. PLOS Computational Biology 20, e1012320. doi:10.1371/journal.pcbi.1012320

Haddix, M.L., Gregorich, E.G., Helgason, B.L., Janzen, H., Ellert, B.H., Francesca Cotrufo, M., 2020. Climate, carbon content, and soil texture control the independent formation and persistence of particulate and mineral-associated organic matter in soil. Geoderma 363, 114160. doi:10.1016/j.geoderma.2019.114160

Hall, E.K., Bernhardt, E.S., Bier, R.L., Bradford, M.A., Boot, C.M., Cotner, J.B., del Giorgio, P.A., Evans, S.E., Graham, E.B., Jones, S.E., Lennon, J.T., Locey, K.J., Nemergut, D., Osborne, B.B., Rocca, J.D., Schimel, J.P., Waldrop, M.P., Wallenstein, M.D., 2018. Understanding how microbiomes influence the systems they inhabit. Nature Microbiology 3, 977–982. doi:10.1038/s41564-018-0201-z

Hansen, P.M., Even, R., King, A.E., Lavallee, J., Schipanski, M., Cotrufo, M.F., 2024. Distinct, direct and climatelJmediated environmental controls on global particulate and minerallJassociated organic carbon storage. Global Change Biology 30, e17080. doi:10.1111/gcb.17080

Hassink, J., 1997. The capacity of soils to preserve organic C and N by their association with clay and silt particles. Plant and Soil 191, 77–87. doi:10.1023/A:1004213929699

Heckman, K.A., Possinger, A.R., Badgley, B.D., Bowman, M.M., Gallo, A.C., Hatten, J.A., Nave, L.E., SanClements, M.D., Swanston, C.W., Weiglein, T.L., Wieder, W.R., Strahm, B.D., 2023. Moisture-driven divergence in mineral-associated soil carbon persistence. Proceedings of the National Academy of Sciences 120, e2210044120. doi:10.1073/pnas.2210044120

Jilling, A., Keiluweit, M., Gutknecht, J.L.M., Grandy, A.S., 2021. Priming mechanisms providing plants and microbes access to mineral-associated organic matter. Soil Biology and Biochemistry 158, 108265. doi:10.1016/j.soilbio.2021.108265

Kaiser, C., Franklin, O., Dieckmann, U., Richter, A., 2014. Microbial community dynamics alleviate stoichiometric constraints during litter decay. Ecology Letters 17, 680–690. doi:10.1111/ele.12269

Kaiser, C., Franklin, O., Richter, A., Dieckmann, U., 2015. Social dynamics within decomposer communities lead to nitrogen retention and organic matter build-up in soils. Nature Communications 6, 8960. doi:10.1038/ncomms9960

Kalinina, O., Cherkinsky, A., Chertov, O., Goryachkin, S., Kurganova, I., Lopes De Gerenyu, V., Lyuri, D., Kuzyakov, Y., Giani, L., 2019. Post-agricultural restoration: Implications for dynamics of soil organic matter pools. CATENA 181, 104096. doi:10.1016/j.catena.2019.104096

Kallenbach, C.M., Frey, S.D., Grandy, A.S., 2016. Direct evidence for microbial-derived soil organic matter formation and its ecophysiological controls. Nature Communications 7, 13630. doi:10.1038/ncomms13630

Kamble, P.N., Bååth, E., 2016. Comparison of fungal and bacterial growth after alleviating induced N-limitation in soil. Soil Biology and Biochemistry 103, 97–105. doi:10.1016/j.soilbio.2016.08.015

King, A.E., Amsili, J.P., Córdova, S.C., Culman, S., Fonte, S.J., Kotcon, J., Liebig, M., Masters, M.D., McVay, K., Olk, D.C., Schipanski, M., Schneider, S.K., Stewart, C.E., Cotrufo, M.F., 2023. A soil matrix capacity index to predict mineral-associated but not particulate organic carbon across a range of climate and soil pH. Biogeochemistry 165, 1–14. doi:10.1007/s10533-023-01066-3

Lavallee, J.M., Soong, J.L., Cotrufo, M.F., 2020. Conceptualizing soil organic matter into particulate and minerallJassociated forms to address global change in the 21st century. Global Change Biology 26, 261–273. doi:10.1111/gcb.14859

Liang, C., Amelung, W., Lehmann, J., Kästner, M., 2019. Quantitative assessment of microbial necromass contribution to soil organic matter. Global Change Biology 25, 3578–3590. doi:10.1111/gcb.14781

Lugato, E., Lavallee, J.M., Haddix, M.L., Panagos, P., Cotrufo, M.F., 2021. Different climate sensitivity of particulate and mineral-associated soil organic matter. Nature Geoscience 14, 295–300. doi:10.1038/s41561-021-00744-x

Malik, A.A., Martiny, J.B.H., Brodie, E.L., Martiny, A.C., Treseder, K.K., Allison, S.D., 2020. Defining trait-based microbial strategies with consequences for soil carbon cycling under climate change. The ISME Journal 14, 1–9. doi:10.1038/s41396-019-0510-0

Manzoni, S., Taylor, P., Richter, A., Porporato, A., Ågren, G.I., 2012. Environmental and stoichiometric controls on microbial carbonlJuse efficiency in soils. New Phytologist 196, 79–91. doi:10.1111/j.1469-8137.2012.04225.x

Mazzilli, S.R., Kemanian, A.R., Ernst, O.R., Jackson, R.B., Piñeiro, G., 2014. Priming of soil organic carbon decomposition induced by corn compared to soybean crops. Soil Biology and Biochemistry 75, 273–281. doi:10.1016/j.soilbio.2014.04.005

Mosier, S., Apfelbaum, S., Byck, P., Calderon, F., Teague, R., Thompson, R., Cotrufo, M.F., 2021. Adaptive multi-paddock grazing enhances soil carbon and nitrogen stocks and stabilization through mineral association in southeastern U.S. grazing lands. Journal of Environmental Management 288, 112409. doi:10.1016/j.jenvman.2021.112409

Poeplau, C., Dechow, R., Begill, N., Don, A., 2024. Towards an ecosystem capacity to stabilise organic carbon in soils. Global Change Biology 30, e17453. doi:10.1111/gcb.17453

Prairie, A.M., King, A.E., Cotrufo, M.F., 2023. Restoring particulate and mineral-associated organic carbon through regenerative agriculture. Proceedings of the National Academy of Sciences 120, e2217481120. doi:10.1073/pnas.2217481120

Rocci, K.S., Cotrufo, M.F., Ernakovich, J., Foster, E., Frey, S., Georgiou, K., Grandy, A.S., Malhotra, A., Reich, P.B., Schlerman, E.P., Wieder, W.R., 2024. Bridging 20 Years of Soil Organic Matter Frameworks: Empirical Support, Model Representation, and Next Steps. Journal of Geophysical Research: Biogeosciences 129, e2023JG007964. doi:10.1029/2023JG007964

Salonen, A.-R., Soinne, H., Creamer, R., Lemola, R., Ruoho, N., Uhlgren, O., De Goede, R., Heinonsalo, J., 2023. Assessing the effect of arable management practices on carbon storage and fractions after 24 years in boreal conditions of Finland. Geoderma Regional 34, e00678. doi:10.1016/j.geodrs.2023.e00678

Sihi, D., Gerber, S., Inglett, P.W., Inglett, K.S., 2016. Comparing models of microbial–substrate interactions and their response to warming. Biogeosciences 13, 1733–1752. doi:10.5194/bg-13-1733-2016

Sistla, S.A., Rastetter, E.B., Schimel, J.P., 2014. Responses of a tundra system to warming using SCAMPS: a stoichiometrically coupled, acclimating microbe–plant–soil model. Ecological Monographs 84, 151–170. doi:10.1890/12-2119.1

Six, J., Conant, R.T., Paul, E.A., Paustian, K., 2002. Stabilization mechanisms of soil organic matter: Implications for C-saturation of soils. Plant and Soil 241, 155–176.

Six, J., Doetterl, S., Laub, M., Müller, C.R., Van De Broek, M., 2024. The six rights of how and when to test for soil C saturation. SOIL 10, 275–279. doi:10.5194/soil-10-275-2024

Smith, P., 2016. Soil carbon sequestration and biochar as negative emission technologies. Global Change Biology 22, 1315–1324. doi:10.1111/gcb.13178

Smith, P., Cotrufo, M.F., Rumpel, C., Paustian, K., Kuikman, P.J., Elliott, J.A., McDowell, R., Griffiths, R.I., Asakawa, S., Bustamante, M., House, J.I., Sobocká, J., Harper, R., Pan, G., West, P.C., Gerber, J.S., Clark, J.M., Adhya, T., Scholes, R.J., Scholes, M.C., 2015. Biogeochemical cycles and biodiversity as key drivers of ecosystem services provided by soils. SOIL 1, 665–685. doi:10.5194/soil-1-665-2015

Stewart, C.E., Paustian, K., Conant, R.T., Plante, A.F., Six, J., 2008. Soil carbon saturation: Evaluation and corroboration by long-term incubations. Soil Biology and Biochemistry 40, 1741–1750. doi:10.1016/j.soilbio.2008.02.014

Stewart, C.E., Paustian, K., Conant, R.T., Plante, A.F., Six, J., 2007. Soil carbon saturation: concept, evidence and evaluation. Biogeochemistry 86, 19–31. doi:10.1007/s10533-007-9140-0

Treseder, K.K., 2008. Nitrogen additions and microbial biomass: a meta-analysis of ecosystem studies. Ecology Letters 11, 1111–1120. doi:10.1111/j.1461-0248.2008.01230.x

Trivedi, P., Anderson, I.C., Singh, B.K., 2013. Microbial modulators of soil carbon storage: integrating genomic and metabolic knowledge for global prediction. Trends in Microbiology 21, 641–651. doi:10.1016/j.tim.2013.09.005

Wang, G., Jagadamma, S., Mayes, M.A., Schadt, C.W., Megan Steinweg, J., Gu, L., Post, W.M., 2015. Microbial dormancy improves development and experimental validation of ecosystem model. The ISME Journal 9, 226–237. doi:10.1038/ismej.2014.120

West, S.A., Diggle, S.P., Buckling, A., Gardner, A., Griffin, A.S., 2007. The Social Lives of Microbes. Annual Review of Ecology, Evolution, and Systematics 38, 53–77. doi:10.1146/annurev.ecolsys.38.091206.095740

West, T.O., Six, J., 2007. Considering the influence of sequestration duration and carbon saturation on estimates of soil carbon capacity. Climatic Change 80, 25–41. doi:10.1007/s10584-006-9173-8

Whalen, E.D., Grandy, A.S., Sokol, N.W., Keiluweit, M., Ernakovich, J., Smith, R.G., Frey, S.D., 2022. Clarifying the evidence for microbiallJ and plantlJderived soil organic matter, and the path toward a more quantitative understanding. Global Change Biology 28, 7167–7185. doi:10.1111/gcb.16413

Wieder, W.R., Grandy, A.S., Kallenbach, C.M., Bonan, G.B., 2014. Integrating microbial physiology and physio-chemical principles in soils with the MIcrobial-MIneral Carbon Stabilization (MIMICS) model. Biogeosciences 11, 3899–3917. doi:10.5194/bg-11-3899-2014

Wu, T., Ost, A.D., Audinot, J.-N., Wiesmeier, M., Wirtz, T., Buegger, F., Häusler, W., Höschen, C., Mueller, C.W., 2022. Association of fresh low-molecular-weight organic compounds with clay-sized mineral fraction in soils of different organic carbon loading. Geoderma 409, 115657. doi:10.1016/j.geoderma.2021.115657

Yang, Y., Dou, Y., Wang, B., Wang, Y., Liang, C., An, S., Soromotin, A., Kuzyakov, Y., 2022. Increasing contribution of microbial residues to soil organic carbon in grassland restoration chronosequence. Soil Biology and Biochemistry 170, 108688. doi:10.1016/j.soilbio.2022.108688

Yu, W., Huang, W., Weintraub-Leff, S.R., Hall, S.J., 2022. Where and why do particulate organic matter (POM) and mineral-associated organic matter (MAOM) differ among diverse soils? Soil Biology and Biochemistry 172, 108756. doi:10.1016/j.soilbio.2022.108756

Zhu, Q., Riley, W.J., Tang, J., Koven, C.D., 2016. Multiple soil nutrient competition between plants, microbes, and mineral surfaces: model development, parameterization, and example applications in several tropical forests. Biogeosciences 13, 341–363. doi:10.5194/bg-13-341-2016

