## Supplemental information for "Microbial community regulation of extracellular enzyme production can mediate patterns of particulate and mineral-associated organic matter accumulation in undersaturated soils"

**Supplementary Table 1.** General model parameters, reproduced following Kaiser et al., 2015 with modifications to match the current model.

| **Parameter** | **Description** | **Value** |
| --- | --- | --- |
| *Enzyme kinetics* | | |
| k_cat_: number of enzymatic reactions catalyzed per enzyme (mol substrate C decomposed mol enzyme C^-1^) | | |
| *k_cat_POM* | k_cat_ of enzymes degrading plant-derived POM | 0.22 timestep^-1*^ |
| *k_cat_CRN* | k_cat_ of enzymes degrading C-rich microbial necromass (i.e., microbial-derived POM) | 0.63 timestep^-1^ |
| *k_cat_NRN* | k_cat_ of enzymes degrading N-rich microbial necromass (i.e., microbial-derived POM) | 0.3 timestep^-1^ |
| k_m_: half saturation constant for substrates in one microsite | | |
| *k_m_POM* | k_m_ of plant-derived POM | 0.29 fmol C |
| *k_m_CRN* | k_m_ of C-rich microbial necromass (i.e., microbial-derived POM) | 0.28 fmol C |
| *k_m_NRN* | k_m_ of N-rich microbial necromass (i.e., microbial-derived POM) | 0.25 fmol C |
| *Microbial physiology* | | |
| *R_maint_* | Maintenance respiration (as fraction of cell biomass) | 0.002667 timestep^-1^ |
| *R_eg_* | Respiration associated with enzyme production and growth (as fraction of C uptake allocated to enzyme production and growth) | 0.26 timestep^-1^ |
| *U_max_* | Maximum C uptake rate (as fraction of cell biomass, to be multiplied by cell surface area:volume ratio) | 0.0019 timestep^-1^ |
| *Pr_inv_* | Probability that an already-occupied microsite will be invaded and colonized by a neighboring, reproducing cell | 0.01 |
| *F_M_* | In the case of random catastrophic death, factor that links maximum cell size to mortality rate | 0.1133 |
| *Initial pools in each microsite* | | |
| *C_enz_* | Active extracellular enzymes | 0.5 fmol C |
| *C_CRN_* | C-rich microbial necromass (i.e., microbial-derived POM) | 100 fmol C |
| *C_NRN_* | N-rich microbial necromass (i.e., microbial-derived POM) | 30 fmol C |
| *C_DOM_* | DOM available for immediate uptake (initial C:N=8)^**^ | 7 fmol C |
| *C_POM_* | Plant-derived POM | 8333 fmol C |
| *Initial number of microsites occupied by microbes* | | |
| *M_O_* | Microsites occupied by microbes, randomly distributed over 10,000 microsites | 1666 |
| ^*^One model timestep=1 h.  ^**^the C:N of immediately assimilable DOM has an initial value of 8, but is not a parameter and can change over the course of the simulation. | | |

**Supplementary Table 2.** Replicate simulations excluded from final analyses due to strongly limiting resource conditions, causing microbial communities to die before they could begin degrading the initial plant-derived particulate organic matter (POM) substrate. Any combinations of cheater presence, initial plant-derived POM C:N, and fr_diff_ not listed in this table were included in downstream analyses with the full ten replicates.

| **Microbial community** | **Initial POM C:N** | ***fr_diff_* value** | **Number of replicates excluded** |
| --- | --- | --- | --- |
| Cheaters + producers | 60 | 0.058 | 1 |
| Cheaters + producers | 70 | 0.0502 | 3 |
| Cheaters + producers | 70 | 0.058 | 1 |
| Cheaters + producers | 80 | 0.0421 | 1 |
| Cheaters + producers | 80 | 0.0502 | 6 |
| Cheaters + producers | 80 | 0.058 | 9 |
| Cheaters + producers | 90 | 0.034 | 1 |
| Cheaters + producers | 90 | 0.0502 | 10 |
| Cheaters + producers | 90 | 0.058 | 9 |
| Cheaters + producers | 100 | 0.0421 | 6 |
| Cheaters + producers | 100 | 0.0502 | 10 |
| Cheaters + producers | 100 | 0.058 | 10 |

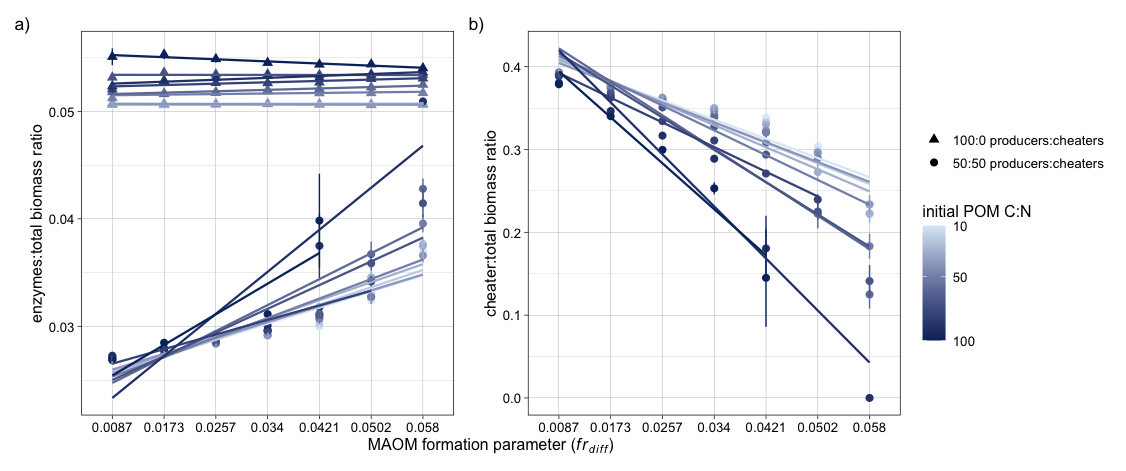

***Supplementary Figure 1*.** Response of a) the ratio of total enzymes produced to total microbial biomass; and b) the ratio of cheater biomass to total microbial biomass to changes in *fr_diff_*. This figure replicates Figure 3 in the main text, with the inclusion of regression lines for easier visualization of variations in the relationships between *fr_diff_* and the above emergent microbial properties with plant-derived particulate organic matter carbon to nitrogen ratio (POM C:N). Shades of blue indicate POM C:N, with darker colors corresponding to higher C:N. Triangles represent simulations that contain only enzyme producers, and circles represent simulations that also contain cheaters. Points represent group means of each initial POM C:N and *fr_diff_* combination, and error bars indicate standard error.

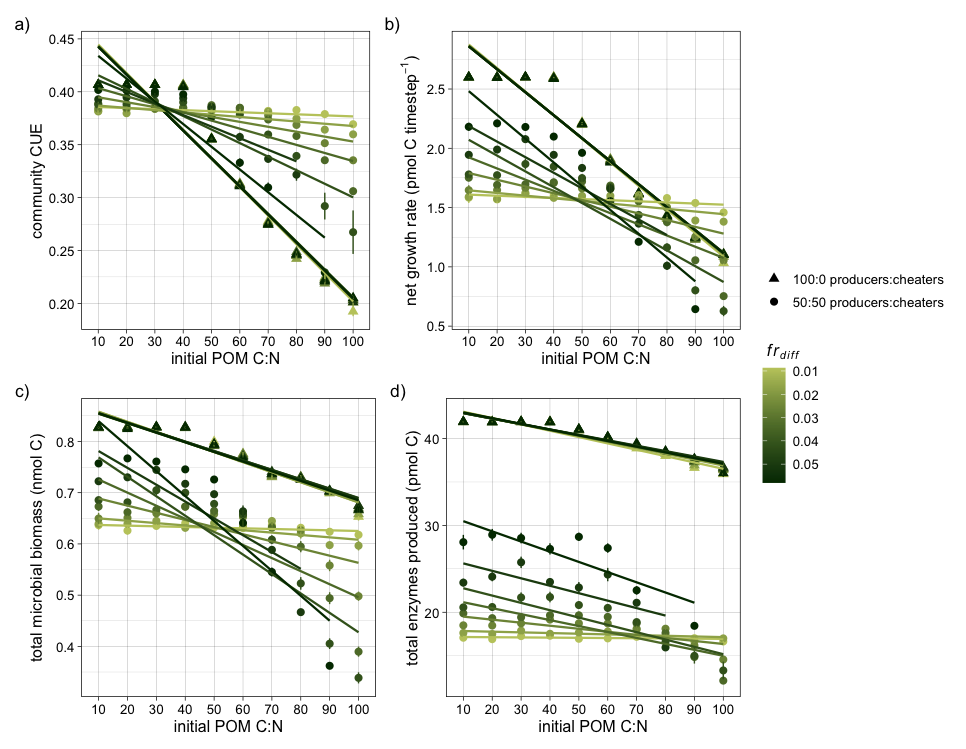

***Supplementary Figure 2*.** Response of a) community carbon use efficiency (CUE); b) net microbial growth rate; c) total microbial biomass; and d) total enzymes produced to initial plant-derived particulate organic matter carbon to nitrogen ratio (POM C:N) and *fr_diff_*. This figure replicates Figure 5 in the main text, with the inclusion of regression lines for easier visualization of the relationships between *fr_diff_* and initial POM C:N. Shades of green indicate values of *fr_diff_*, with darker shades corresponding to higher *fr_diff_*. Triangles correspond to simulations containing only enzyme producers, and circles to simulations that also contain cheaters. All simulations begin with the same microbial biomass, and simulations that contain cheaters begin with a 50:50 mix of enzyme producers and cheaters. Points represent group means of each initial plant-derived POM C:N and *fr_diff_* combination, and error bars indicate standard error.

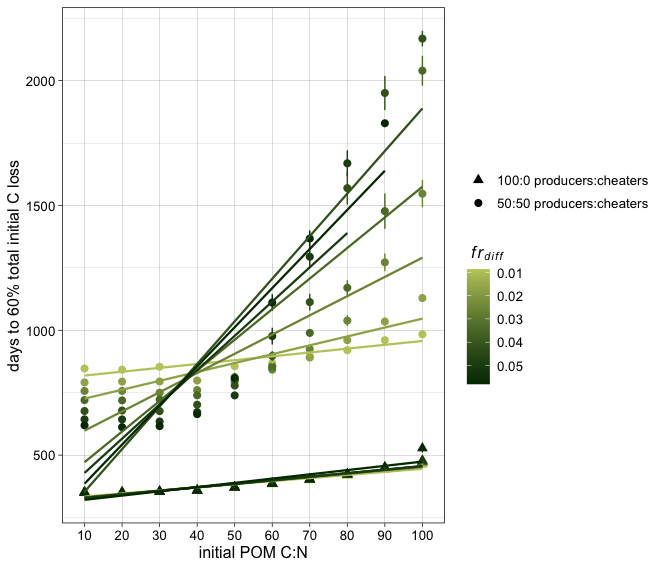

***Supplementary Figure 3*.** The amount of time it takes the microbial community to degrade 60% of the total initial carbon (C) pool in response to initial plant-derived particulate organic matter carbon to nitrogen ratio (POM C:N) and *fr_diff_*. This figure replicates Figure 5 in the main text, with the inclusion of regression lines for easier visualization of the relationships between *fr_diff_* and initial POM C:N. Shades of green indicate values of *fr_diff_*, with darker shades corresponding to higher *fr_diff_*. Triangles represent simulations that contain only enzyme producers, and circles represent those that also contain cheaters. Points represent group means of each plant-derived POM C:N and *fr_diff_* combination, and error bars indicate standard error.

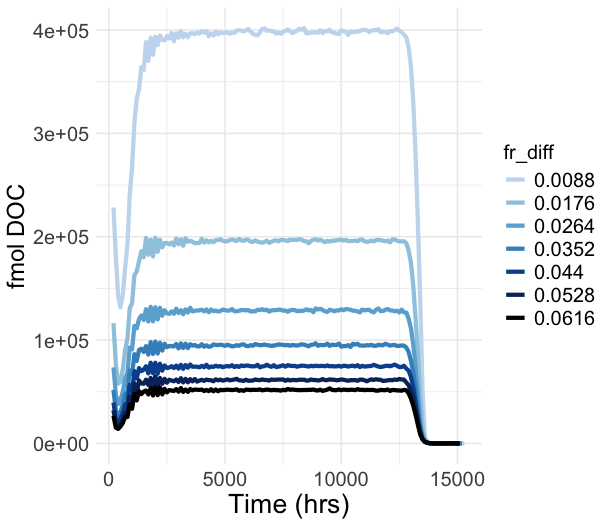

***Supplementary Figure 4*.** Time-series plot of the total amount of dissolved organic carbon (DOC) available on the grid to microbes throughout the course of simulations, when initial particulate organic matter carbon to nitrogen ratio (POM C:N) is 10. Different shades of blue indicate values *fr_diff_* (i.e., MAOM saturation state).
